# OMEinfo: Global Geographic Metadata for -omics Experiments

**DOI:** 10.1101/2023.10.23.563576

**Authors:** Matthew Crown, Matthew Bashton

## Abstract

Microbiome studies increasingly associate geographical features like rurality and climate types with microbiomes. However, microbiologists/bioinformaticians often struggle to access and integrate rich geographical metadata from sources such as GeoTIFFs; and inconsistent definitions of rurality, for example, can hinder cross-study comparisons. To address this, we present OMEinfo, a Python-based tool for automated retrieval of consistent geographical metadata from user-provided location data. OMEinfo leverages open data sources such as the Global Human Settlement Layer, Köppen-Geiger climate classification models, and Open-Data Inventory for Anthropogenic Carbon dioxide, to ensure metadata accuracy and provenance. OMEinfo’s Dash application enables users to visualise their sample metadata on an interactive map and to investigate the spatial distribution of metadata features, which is complemented by data visualisation to analyse patterns and trends in the geographical data before further analysis. The tool is available as a Docker container, providing a portable, lightweight solution for researchers. Through its standardised metadata retrieval approach and incorporation of FAIR and Open data principles, OMEinfo promotes reproducibility and consistency in microbiome metadata. To demonstrate its utility, OMEinfo is utilised to replicate the results of a previous study linking population density to soil sample alpha diversity. As the field continues to explore the relationship between microbiomes and geographical features, tools like OMEinfo will prove vital in developing a robust, accurate, and interconnected understanding of these interactions, whilst having applicability beyond this field to any studies utilising location-based metadata. Finally, we release the OMEinfo annotation dataset, a collection of 5.3 million OMEinfo annotated samples from the ENA, for use in a retrospective analysis of sequencing samples, and highlight a number of ways researchers and sequencing read repositories can improve the quality of underlying metadata submitted to these public stores.

**Availability:** OMEinfo is freely available and released under an MIT licence. OMEinfo source code is available at https://github.com/m-crown/OMEinfo/

## 1. Introduction

The microbial ecology of the built environment plays a role in shaping our human health[1], [2], the environment around us, *e.g*. through microbially induced corrosion[3] and can be harnessed for bioremediation, such as in oil spill clean up[4], and with activated sludge in wastewater treatment plants[5]. Recently, much focus has been placed on associating geographical features such as rurality[6], elevation[7], climate type, and even specific cities[8] to microbiomes. For example, Wang *et al*. demonstrate a positive correlation between population and soil bacterial diversity[9], whilst urbanisation has been demonstrated to alter the soil rhizosphere microbiome across a gradient from urban to rural[10]. As the evidence for such associations has grown, so has the size of studies being performed to examine these effects[8], [11], [12]. With global-scale studies, comes a need for globally consistent metadata annotation, for both consistency within a study, as well as for metanalysis of global studies.

These projects require the determination of geographical metadata features, typically through online data sources such as Google Maps for latitude, longitude and elevation or GIS services such as ArcGIS Online. Microbiologists/bioinformaticians are typically not familiar with the plethora of geographical data available publicly through *e.g*. satellite-derived GeoTIFFs, and even so, may not be aware of the tools and methods required to integrate these into their analysis. This has led to many studies in the field employing an ambiguous, approximate or sub-optimal definition of many environmental characteristics, including rurality[2], [6], [10], [13]–[15] and climate classification[16], limiting the ability to compare between studies and re-analyse data in a global context (see Table S1 for a more comprehensive list of differences between study definitions of such metadata features and their issues).

In addition, some measurements, such as rurality, are often determined manually, either by an assumed value for a location or through a determination made by local authorities or governments. This can lead to issues when comparing across studies, where definitions may vary significantly - *i.e*. what is considered rural in a densely populated country may be considered urban in another country (see Table 1).

**Table 1.** A sample of the different definitions of rurality in use across the world.

| Country | Urban/Rural Definition | Reference |
| --- | --- | --- |
| Canada | An urban area was defined as having a population of at least 1,000 and a density of 400 or more people per square kilometre | [17] |
| USA | ~2500 inhabitants/km<sup>2</sup>(1000/mile<sup>2</sup>) | [18] |
| Japan | Define “Densely Inhabited Districts” as those with 4000+ inhabitants/km<sup>2</sup> | [19] |
| UK | Rural defined as areas which “fall outside of settlements with more than 10,000 resident population” | [20] |
| Iceland | “Rural areas are agricultural areas or villages with less than 200 inhabitants” | [21] |
| Norway | Urban defined as minimum 200 inhabitants per cluster of buildings. Minimum population density in urban settlements (2022) = 236/km<sup>2</sup> | [22] |

To be consistent in the use of geographical metadata, it is therefore crucial that the source of metadata features are communicated, something which is often overlooked, and where possible, globally consistent definitions for these features are used. To date, no tool exists to provide such geographical/cultural/socioeconomic metadata feature annotation in a globally consistent manner. To this end, we present OMEinfo, a tool for the automated retrieval of consistent geographical metadata including Köppen-Geiger climate classification, degree of rurality, population density, CO_2_ and NO_2_ emissions and relative deprivation from user-provided location data, together with a versioned data source which allows for metadata provenance.

## 2. Implementation

OMEinfo is released as a Dash based web app, designed to run locally in the browser. An overview of the OMEinfo workflow can be seen in Fig. 1.

**Fig. 1.**
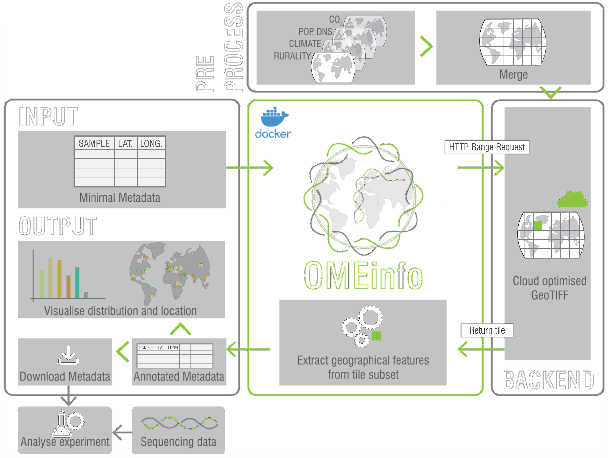
The OMEinfo workflow. Users upload a metadata file with sample, latitude and longitude to the Dockerised web app (or CLI). OMEinfo processes locations and retrieves relevant sub-tiles from the Cloud Optimised GeoTIFF necessary for annotating the location. Features are annotated for each location and upon completion, interactive maps and plots are displayed to the user. Metadata and citations can then be downloaded and integrated into downstream analysis.

### 2.1. Data Aggregation and Preprocessing

OMEinfo utilises cloud-optimised GeoTIFFs (COGs) for back-end data storage, hosted online in an AWS S3 bucket. Various data sources are integrated into the OMEinfo v2 dataset covering geographical, environmental and socioeconomic characteristics (see Table S2).

All data sources are reprojected into World Geodetic System 1984 (WGS84/ESRI:4326) format and merged into a single multi-band file using Geospatial Data Abstraction Library (GDAL) v3.7.0[23]. The multi-band WGS84 projection file is then converted to COG format using Rasterio v1.3.7[24] and the rio-cogeo v5.0 plugin[25]. This COG file is then uploaded to an AWS S3 bucket, and is queried by the OMEinfo web-app (see below), but can also be downloaded by end-users should they wish to interact directly with the data sources utilised in OMEinfo.

### 2.2. OMEinfo

OMEinfo is available as both a Dockerised web-app (OMEinfo app) and a command-line tool (OMEinfo CLI). OMEinfo app is built with Dash v2.9.3[26] and Dash-Bootstrap v1.4.1[27]. OMEinfo CLI provides a convenient method for power users to process data directly in the command-line, allowing for batch processing and integration into pipelines, and additionally makes use of Textualize Rich[28] for rich text output in the terminal. Both the CLI and app require users to upload a metadata sheet including sample name, latitude and longitude in CSV/TSV format. Latitude and longitude should be in the common EPSG:4326 coordinate system, which is a format used by all common web-based mapping tools e.g. Google Maps and Open Street Map.

OMEinfo utilises rio-cogeo, Rasterio and GDAL to query the COG files stored in the OMEinfo S3 bucket. Queries take the form of HTTPS range requests, which return small, but not uniquely identifiable, subsets of the OMEinfo COG locally, where they are queried for specific point data. GDAL supports query caching, meaning that if locations are geographically similar, locally cached data storage can significantly speed up annotation.

OMEinfo has been designed with data privacy in mind. By using GDAL to submit range requests, rather than transmitting the locations themselves, OMEinfo keeps sensitive location metadata private, helping to keep personally identifiable information secure, and allowing for metadata to be distributed with globally consistent metadata features, whilst maintaining anonymisation of data. This feature benefits studies in which ethical approvals prevent the sharing of location data, for example, sampling in an individual’s home. Following the completion of the analysis, OMEinfo app presents users with a summary of annotated metadata fields, in the form of interactive histograms, maps and tables, built using Plotly Express[29] which allows users to see the OMEinfo metadata in context.

In addition to the analysis, OMEinfo app includes an explore page that allows users to interact with a 3D digital elevation model (DEM) map representation of the underlying data sources, built using QGIS[30] and Qgis2threejs[31] to aid an understanding of the structure of the underlying data without needing to download any additional files.

OMEinfo is committed to the principles of FAIR data sharing:

- Findable: All code and data sources essential for constructing OMEinfo data packets are openly accessible within the repository.
- Accessible: To foster accessibility, all OMEinfo code is released under the permissive MIT licence, and the GeoTIFF data utilised within OMEinfo adheres to permissive data usage licenses, as specified in Table S2.
- Interoperable: OMEinfo prioritises interoperability by citing all data sources employed within the application in a downloadable BibTex format, both within the web application and the repository.
- Reusable: By versioning the formatted data packets upon release and documenting their contents in a changelog within the repository, OMEinfo ensures that previous data versions remain accessible for continued use, encouraging data reusability and promoting sustainability in research efforts.

## 3. Applications and Examples

### 3.1. Potential Use Cases

OMEinfo is usable in any study employing sampling which can be associated with a specific geographic location. It allows for automated and consistent metadata annotation based on these locations. The tool was originally developed for use with the built environment metagenomics analysis (or any -omics data, hence OME) with potential use cases including:

- Determining associations between microbiome diversity and geographical metadata features, *e.g*. population density (section 3.2).
- As a ground truth dataset for classification algorithms utilising microbial taxa abundances for location classification.
- To identify locations with specific environmental conditions suitable for microbial biotechnological applications (*e.g*., bioprospecting for extremophiles in areas with high NO_2_ concentrations).

### 3.2. Correlation between OMEinfo-derived and census-derived metadata

Wang *et al*. utilise population density estimates derived from US census data and census tract blocks to determine a population density/km^2[9]^. To verify the accuracy of population density annotation from OMEinfo (in v2 data packet at 1km^2^ resolution), Spearman’s rank correlation analysis was used to assess the relationship between OMEinfo population densities and the original study’s census-tract defined population densities. The results revealed a strong positive and statistically significant correlation (Spearman’s ρ = 0.78, p < 0.01).

Some variability between census-tract and OMEinfo-derived values is observed at higher population densities (Fig. S2). This is likely due to the difference in method for estimating population density: in the Global Human Settlement Layer Population (GHSL-POP) GeoTIFF, a population disaggregation model is used to distribute the underlying population to built-up areas, whereas a census is able to precisely determine the population living in a given area. However, the benefit of the OMEinfo approach to assigning population density still remains, in that from a single web app/CLI tool, population density, together with a host of additional metadata features, can be determined without needing to obtain individual census data, making it well suited to providing consistent metadata for complex global studies.

To verify the applicability of OMEinfo in a biological context, Spearman’s rank correlation was utilised to re-evaluate the finding previously reported in the same study, in which population density showed a positive association with Shannon diversity and phylogenetic diversity.

A positive relationship was observed between diversity and population density, using OMEinfo-derived population density and Census Tract population density. Shannon diversity was positively correlated with Census Tract and OMEinfo Population Density with coefficients of 0.225 and 0.214, respectively (p < 0.01). Similarly, Phylogenetic Diversity exhibited a correlation of 0.259 with Census Tract Population Density and 0.244 with OMEinfo Population Density, both statistically significant (p < 0.01).

### 3.3. OMEinfo annotation dataset

The European Nucleotide Archive (ENA)[32] sequence read archive serves as a central repository of read data. As part of submission to ENA, users may optionally choose to distribute location data for associated biosamples. From a total of 34,716,112 samples queried [access date: 2023-10-13], 5,361,056 samples had an associated latitude and longitude making them suitable for processing with OMEinfo. This represents an ideal large-scale dataset for benchmarking the performance of OMEinfo, as well as providing an opportunity to distribute a precomputed OMEinfo dataset well suited to meta-analysis of existing studies which further improves the availability of globally consistent geographic metadata.

Through analysis with OMEinfo CLI v1 and a local copy of the OMEinfo v2 dataset, we were able to annotate the metadata of 5,338,430 samples in ENA with geographical metadata (22,626 samples contained invalid latitude/longitude data) in less than 30 minutes single-threaded (Intel Xeon Gold 6230 CPU @ 2.10GHz, Rocky Linux release 8.8). An additional dataset was produced covering samples specifically marked as environmental (metadata feature describes samples derived from an environmental DNA sample) in ENA. A total of 190,573 samples are marked as environmental, with 135,517 containing valid coordinates for analysis. The total analysis time for this dataset (same parameters as full) took 60.27 seconds. These annotation files can be downloaded from the S3 Bucket (see GitHub repo for more info).

We find that environmental samples in the ENA show a skew towards the extremes of the urban-rural scale (Fig. 2A), with 78.59% of all environmental samples annotated as urban centres (40,565 samples), very low density (39,716 samples) or water (26,216 samples). Future global studies of rurality/urbanisation should ensure that focus is given to all levels of the urban-rural continuum, as it is possible that additional insights could be gained by sampling from understudied areas of such a continuum.

**Fig. 2.**
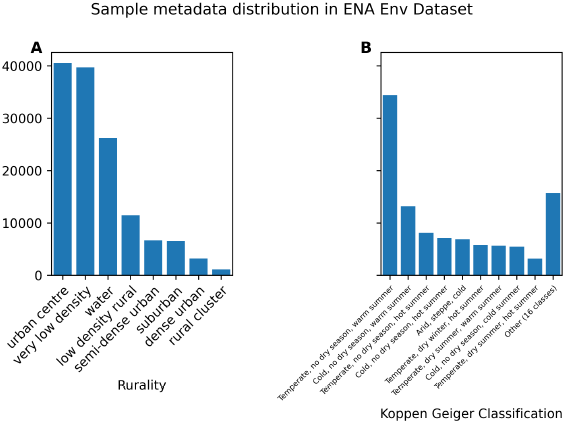
Sample metadata distribution in the ENA Env. dataset. A: The breakdown of samples according to the rurality index shows the majority of samples are from urban centres, very low-density locations or water. B: Breakdown of samples according to Köppen-Geiger Climate classification. Most samples are taken from “Temperate, no dry season, warm summer” climates.

This pattern is even more pronounced for Köppen-Geiger climate classifications, where 25.41% of all environmental samples are from locations with a “*Temperate, no dry season, warm summer*” climate type (Fig. 2B). This is especially relevant as multiple studies have shown that climate type is a driver of microbiome diversity in the built-environment[8] as well as other ecosystems including streams[33] and on stone surfaces of cultural heritage sites[34].

Within the full ENA dataset, more than 3,700 samples report a location of 0.00N 0.00E for their sample, a location in the South Atlantic Ocean. Samples with this location also have “Environment Biome” metadata values including the following:

- Wetland
- Human-gut
- International Space Station - Russian Service Module
- Anthropogenic forest
- Large river biome

From this, it is clear that certain locations, including 0.00N 0.00E, are used as placeholders to satisfy metadata requirements. Within the environmental database, we can see a number of frequent sample locations occurring. Whilst some of these represent genuine sample locations, *e.g*. a site in De Mossel (NL), where long-term experiments have been occurring, many others misrepresent the location reported in studies themselves, for example, 1,341 samples from SARS-CoV-2 wastewater sequencing in Liverpool (UK) have a reported location in the North Sea.

To this end, we provide a set of recommendations for submitters to ENA of additional best practices, on top of following existing metadata standards, for submitting data to public read repositories in order to provide the greatest value to downstream users of this data:

- Avoid use of pseudo latitude and longitudes for samples where this data is not available.
- Submit locations at the maximum possible resolution allowable by ethical approvals. In most cases, data can be collected at a high resolution using a modern smartphone, for example with Google Maps or What3Words, and easily converted to latitude and longitude for upload.
- Where sample locations are not able to be shared, use International Nucleotide Sequence Database Collaboration (INSDC)-recommended controlled vocabulary to report this, *e.g*. “not applicable”, “not collected”, “not provided” or “restricted access”[35].

Further, whilst public read repositories such as the ENA have made encouraging updates to minimum metadata standards, including requiring the country of collection for all new samples[36], implementing additional checks for consistency between metadata fields upon submission could ensure the quality of data stored is not avoidably diminished, particularly improving the geographical remediation of metadata records. For example, with these new collection country metadata requirements, it is possible to determine whether coordinates correlate with the reported location and flag this to the user. Additionally, it is possible with OMEinfo to determine if sample coordinates are on a body of water, which can then be sanity-checked against reported “Environment Biome” metadata fields.

Finally, we observed that 22,626 samples had invalid latitude and longitude metadata fields within the ENA. With the exception of checklists ERC000028 and ERC000011 (the ENA default sample checklist), where lat_lon is an optional free text field, all location fields are in the WGS84 decimal degrees format, with submission controlled by the following regular expression:

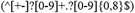

In WGS84 decimal degree format, latitudes should be between ±180 degrees and longitudes between ±90 degrees. Valid regular expressions for this regular expression include 12345 and 1234567890a12345678, and we observe latitude and longitudes orders of magnitude greater than possible in WGS84 coordinate system. We recommend consolidation of all location-based checklist fields to the WGS84 decimal degree format, and validation of submitted metadata to ensure that latitudes and longitudes fall within the expected bounds of the coordinate system. Such remediation will help to improve the quality of submitted data and ensure communication of geolocation is clear and consistent for samples in the ENA regardless of their chosen checklist.

## 4. Conclusion

We present a powerful, innovative and FAIR tool for annotating metadata with essential geographical features. This solution not only enhances the granularity of microbial sequencing metadata but also opens up new possibilities for understanding the interplay between microorganisms and their environments, whilst having broad applicability beyond microbial ecology to any dataset with location-based features. To enable researchers to begin using such data, we present “OMEinfo ENA” and “OMEinfo ENA Env.” datasets; metadata annotations of all samples in the ENA with reported locations, as well as the OMEinfo tool in a pre-built Docker container and a simple CLI tool.

## Availability Statement

OMEinfo is freely available and released under an MIT licence. OMEinfo source code is available at **https://github.com/m-crown/OMEinfo/**.

## Acknowledgements

The authors express their sincere gratitude to Layla van Ellen for developing the OMEinfo logo and assisting in producing Fig 1, and to Dr Haitao Wang for the data necessary for comparing OMEinfo to existing methods.

## References

[1] B. G. Wu et al., “Evidence for Environmental-Human Microbiota Transfer at a Manufacturing Facility with Novel Work-related Respiratory Disease,” Am. J. Respir. Crit. Care Med., vol. 202, no. 12, pp. 1678–1688, Dec. 2020.

[2] P. V. Kirjavainen et al., “Farm-like indoor microbiota in non-farm homes protects children from asthma development,” Nat. Med., vol. 25, no. 7, pp. 1089–1095, Jul. 2019.

[3] Salgar-Chaparro Silvia J., Lepkova Katerina, Pojtanabuntoeng Thunyaluk, Darwin Adam, and Machuca Laura L., “Nutrient Level Determines Biofilm Characteristics and Subsequent Impact on Microbial Corrosion and Biocide Effectiveness,” Appl. Environ. Microbiol., vol. 86, no. 7, pp. e02885–19, Mar. 2020.

[4] A. Sherry et al., “Anaerobic biodegradation of crude oil under sulphate-reducing conditions leads to only modest enrichment of recognized sulphate-reducing taxa,” Int. Biodeterior. Biodegradation, vol. 81, pp. 105–113, Jul. 2013.

[5] S. Begmatov et al., “The structure of microbial communities of activated sludge of large-scale wastewater treatment plants in the city of Moscow,” Sci. Rep., vol. 12, no. 1, p. 3458, Mar. 2022.

[6] A. Parajuli et al., “Urbanization Reduces Transfer of Diverse Environmental Microbiota Indoors,” Front. Microbiol., vol. 9, p. 84, Feb. 2018.

[7] M. Delgado-Baquerizo et al., “Global homogenization of the structure and function in the soil microbiome of urban greenspaces,” Sci Adv, vol. 7, no. 28, p. eabg5809, Jul. 2021.

[8] D. Danko et al., “A global metagenomic map of urban microbiomes and antimicrobial resistance,” Cell, vol. 184, no. 13, pp. 3376–3393.e17, Jun. 2021.

[9] H. Wang, M. Cheng, M. Dsouza, P. Weisenhorn, T. Zheng, and J. A. Gilbert, “Soil Bacterial Diversity Is Associated with Human Population Density in Urban Greenspaces,” Environ. Sci. Technol., vol. 52, no. 9, pp. 5115–5124, May 2018.

[10] C. L. Rosier, S. W. Polson, V. D’Amico 3rd, J. Kan, and T. L. E. Trammell, “Urbanization pressures alter tree rhizosphere microbiomes,” Sci. Rep., vol. 11, no. 1, p. 9447, May 2021.

[11] A. Barberán et al., “Continental-scale distributions of dust-associated bacteria and fungi,” Proc. Natl. Acad. Sci. U. S. A., vol. 112, no. 18, pp. 5756–5761, May 2015.

[12] X. Fu et al., “Continental-Scale Microbiome Study Reveals Different Environmental Characteristics Determining Microbial Richness, Composition, and Quantity in Hotel Rooms,” mSystems, vol. 5, no. 3, May 2020, doi: 10.1128/mSystems.00119-20.

[13] R. I. Adams et al., “Microbial exposures in moisture-damaged schools and associations with respiratory symptoms in students: A multi-country environmental exposure study,” Indoor Air, vol. 31, no. 6, pp. 1952–1966, Nov. 2021.

[14] J. Riedler et al., “Exposure to farming in early life and development of asthma and allergy: a cross-sectional survey,” Lancet, vol. 358, no. 9288, pp. 1129–1133, Oct. 2001.

[15] R. Gacesa et al., “Environmental factors shaping the gut microbiome in a Dutch population,” Nature, vol. 604, no. 7907, pp. 732–739, Apr. 2022.

[16] J. Chase et al., “Geography and Location Are the Primary Drivers of Office Microbiome Composition,” mSystems, vol. 1, no. 2, pp. mSystems.00022–16, e00022–16, Apr. 2016.

[17] Government of Canada and S. Canada, “Population Centre and Rural Area Classification 2016 - Definitions,” Jun. 27, 2018. https://www.statcan.gc.ca/en/subjects/standard/pcrac/2016/definitions (accessed Oct. 13, 2023).

[18] J. Cromartie, “What is Rural?” https://www.ers.usda.gov/topics/rural-economy-population/rural-classifications/what-is-rural/ (accessed Oct. 13, 2023).

[19] Statistics Bureau, Ministry of Internal Affairs, and Communications, “What is a Densely Inhabited District?” https://www.stat.go.jp/english/data/chiri/did/1-1.html (accessed Oct. 13, 2023).

[20] UK Government, “Defining Rural Areas.” Mar. 2017. [Online]. Available: https://assets.publishing.service.gov.uk/government/uploads/system/uploads/attachment_data/file/597751/Defining_rural_areas__Mar_2017_.pdf

[21] “Statistics Iceland: Fewer than 6% of the population live in rural areas,” Statistics Iceland. https://statice.is/publications/news-archive/inhabitants/population-by-urban-nuclei-2001-2020/ (accessed Oct. 13, 2023).

[22] “Population and land area in urban settlements,” SSB. https://www.ssb.no/en/befolkning/folketall/statistikk/tettsteders-befolkning-og-areal (accessed Oct. 13, 2023).

[23] GDAL/OGR contributors, “GDAL/OGR Geospatial Data Abstraction software Library.” Open Source Geospatial Foundation, 2023. doi: 10.5281/zenodo.5884351.

[24] S. Gillies and Others, Rasterio: geospatial raster I/O for Python programmers. Mapbox, 2013-. [Online]. Available: https://github.com/rasterio/rasterio

[25] rio-cogeo: Cloud Optimized GeoTIFF creation and validation plugin for rasterio. Github. Accessed: Oct. 13, 2023. [Online]. Available: https://github.com/cogeotiff/rio-cogeo

[26] dash: Data Apps & Dashboards for Python. No JavaScript Required. Github. Accessed: Oct. 13, 2023. [Online]. Available: https://github.com/plotly/dash

[27] dash-bootstrap-components: Bootstrap components for Plotly Dash. Github. Accessed: Oct. 13, 2023. [Online]. Available: https://github.com/facultyai/dash-bootstrap-components

[28] rich: Rich is a Python library for rich text and beautiful formatting in the terminal. Github. Accessed: Oct. 13, 2023. [Online]. Available: https://github.com/Textualize/rich

[29] plotly_express: Plotly Express - Simple syntax for complex charts. Now integrated into plotly.py! Github. Accessed: Oct. 13, 2023. [Online]. Available: https://github.com/plotly/plotly_express

[30] QGIS Development Team, “QGIS Geographic Information System,” QGIS Association. [Online]. Available: https://www.qgis.org

[31] M. Akagi, Qgis2threejs: A QGIS plugin to export 3D maps to Web. Github. Accessed: Oct. 13, 2023. [Online]. Available: https://github.com/minorua/Qgis2threejs

[32] J. Burgin et al., “The European Nucleotide Archive in 2022,” Nucleic Acids Res., vol. 51, no. D1, pp. D121–D125, Jan. 2023.

[33] F. Picazo et al., “Climate mediates continental scale patterns of stream microbial functional diversity,” Microbiome, vol. 8, no. 1, p. 92, Jun. 2020.

[34] X. Ding, W. Lan, A. Yan, Y. Li, Y. Katayama, and J.-D. Gu, “Microbiome characteristics and the key biochemical reactions identified on stone world cultural heritage under different climate conditions,” J. Environ. Manage., vol. 302, no. Pt A, p. 114041, Jan. 2022.

[35] International Nucleotide Sequence Database Collaboration, “Missing Value Reporting - INSDC.” https://www.insdc.org/submitting-standards/missing-value-reporting/(accessed Oct. 19, 2023).

[36] International Nucleotide Sequence Database Collaboration, “INSDC Spatiotemporal metadata minimum standards update 03-03-2023.” https://www.insdc.org/news/insdc-spatiotemporal-metadata-minimum-standards-update-03-03-2023/ (accessed Oct. 19, 2023).

[37] R. R. Dunn, N. Fierer, J. B. Henley, J. W. Leff, and H. L. Menninger, “Home life: factors structuring the bacterial diversity found within and between homes,” PLoS One, vol. 8, no. 5, p. e64133, May 2013.

[38] Joint Research Centre (European Commission) et al., GHSL data package 2019: public release GHS P2019. LU: Publications Office of the European Union, 2019.

[39] T. Oda and S. Maksyutov, “ODIAC Fossil Fuel CO2 Emissions Dataset,” Center for Global Environmental Research, National Institute for Environmental Studies, 2015, doi: 10.17595/20170411.001.

[40] F. Romahn, M. Pedergnana, D. Loyola, A. Apituley, M. Sneep, and J. P. Veefkind, “Sentinel-5 Precursor/TROPOMI Level 2 Product User Manual: Cloud Properties,” European Space Agency (ESA), 2022.

[41] H. Beck, N. Zimmermann, T. McVicar, N. Vergopolan, A. Berg, and E. Wood, “Present and future Köppen-Geiger climate classification maps at 1-km resolution,” Scientific Data, vol. 5, p. 180214, 2018.

[42] Center for International Earth Science Information Network - CIESIN - Columbia University, “Global Gridded Relative Deprivation Index (GRDI), Version 1.” NASA Socioeconomic Data and Applications Center (SEDAC), Palisades, New York, 2022. doi: 10.7927/3xxe-ap9

