## Supplementary Data for "OMEinfo: Global Geographic Metadata for -omics Experiments"

**Table S1.** Varying sources of metadata in existing microbial ecology studies.

| Study | Geospatial Info (Source) | Metadata Feature Issue | Reference |
|---|---|---|---|
| Geography and Location Are the Primary Drivers of Office Microbiome Composition | Köppen-Geiger climate classification (Unspecified) | Source of climate classification unspecified. | [1] |
| Urbanization pressures alter tree rhizosphere microbiomes | Rurality/Urbanisation (US Census Bureau) | Uses a national-level definition of rurality. | [2] |
| Continental-scale distributions of dust-associated bacteria and fungi | Location | 1-degree resolution latitude/longitude - approximately 11.1km resolution. | [3] |
| Farm-like indoor microbiota in non-farm homes protects children from asthma development | Rurality and location | Rurality reported without methodology. Latitude/longitude is only available on request. | [4] |
| Home Life: Factors structuring the bacterial diversity found within and between homes | Location | Location is constant for 40 homes in a 1000km$^2$ region. | [5] |
| Microbial exposures in moisture-damaged schools and associations with respiratory symptoms in students: A multi-country environmental exposure study | Rurality and location | Building setting reported but the method used is not described. Sample location reported as nearest city, with no latitude or longitude. | [6] |
| Urbanization Reduces Transfer of Diverse Environmental Microbiota Indoors | Rurality and location | Constant latitude/longitude reported for 26 urban and 30 rural homes. Rurality uses a national-level definition. | [7] |
| Exposure to farming in early life and development of asthma and allergy: a cross-sectional survey | Rurality | Population density and farming characteristics considered for region similarity, but data sources and methodology not described. | [8] |
| Environmental factors shaping the gut microbiome in a Dutch population | Rurality (Statistics Netherlands) | Uses a national-level definition of rurality. | [9] |

**Table S2.** OMEinfo v2 data sources

| Annotation | Source | Licence | Reference |
|---|---|---|---|
| Rurality and Population Density | Global Human Settlement Layer | CC BY 4.0 | [10] |
| Fossil Fuel $CO_2$ Emissions | ODIAC | CC BY 4.0 | [11] |
| Tropospheric $NO_2$ Emissions | Copernicus Sentinel S5p | (see SX) | [12] |
| Köppen-Geiger Climate Classification | Beck et al. | CC BY 4.0 | [13] |
| Relative Deprivation | SEDAC | CC BY 4.0 | [14] |

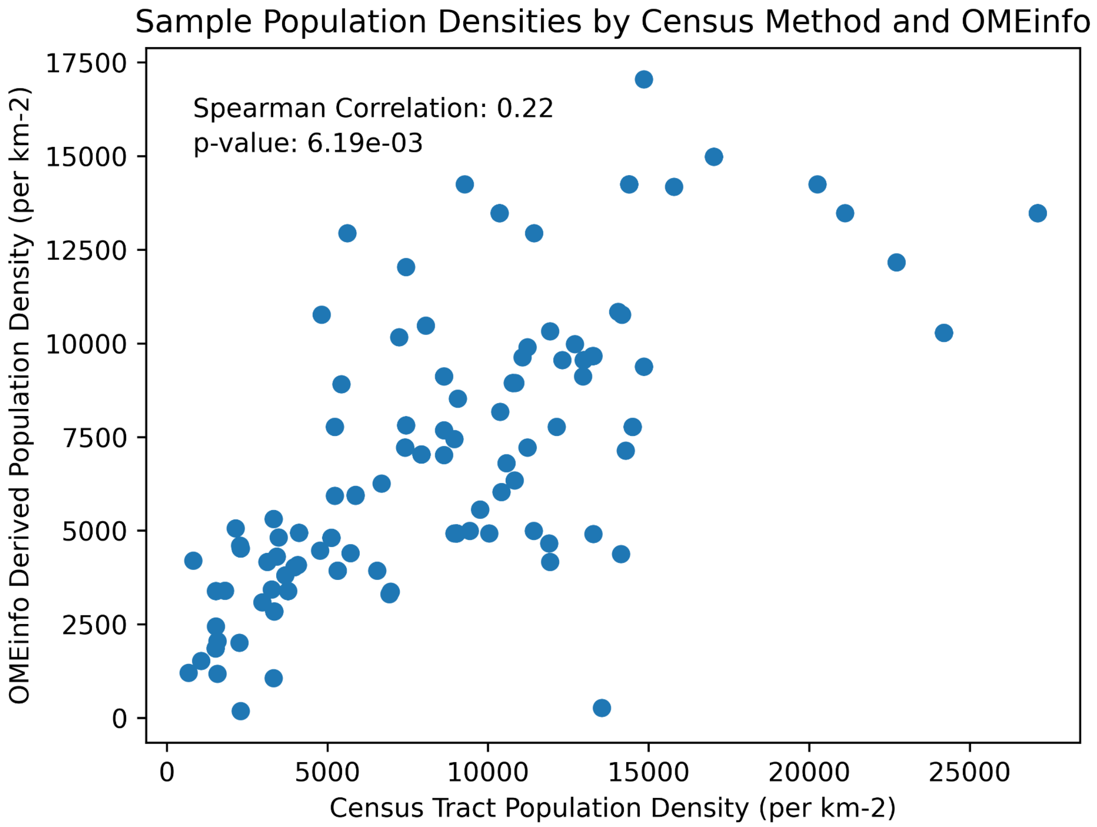


**Fig. S2.**: Correlation between census-tract derived and OMEinfo derived population densities.

### References


[1] J. Chase et al., “Geography and Location Are the Primary Drivers of Office Microbiome Composition,” mSystems, vol. 1, no. 2, pp. mSystems.00022–16, e00022–16, Apr. 2016.
